# Potential Utility of Routine Programmatic Data in Monitoring National and State-Level HIV Epidemic in Nigeria: Data Triangulation Analysis

**DOI:** 10.1101/734293

**Authors:** Debem Henry, Aminu Yakubu, Mukhtar Ahmed, Gwamna Jerry, Dalhatu Ibrahim

## Abstract

Nigeria relies on data from periodic resource-intensive surveys such as antenatal HIV seroprevalence sentinel surveys (ANC-HSS) and population-based National AIDS and Reproductive Health Surveys (NARHS) for its HIV control efforts. Nigeria has not explored the use of readily available routine programmatic data (RPD) to easily inform and monitor epidemic control efforts at local settings in near real time. This study aimed to determine the utility of RPDs (Prevention of Mother-To-Child Transmission [PMTCT] and HIV Testing and Counseling [HTC]) as a proxy for monitoring HIV epidemic in Nigeria. Using World Health Organization 12 step triangulation procedures, we compared state-level seropositivity data from PMTCT and HTC programs to HIV prevalence data from NARHS and ANC-HSS reports in relevant pairs from 2010 to 2014 in Nigeria. The study population was pregnant women and general population. We abstracted relevant data from PEPFAR Nigeria data source and published national survey reports. We compared visual (scatterplots and maps) patterns and trends, and performed Pearson correlation and univariate linear regression models of the estimates for best matched/contiguous years for which data were available. Correlation between PMTCT2014 and ANC-HSS2014 was positive and significant (R=0.7,p<0.001). ANC-HSS2014 and HTC2014 were slightly correlated (R=0.4,p<0.05). Significant correlation was observed between ANC-HSS2010 and PMTCT2013 (R=0.8,p<0.001) and between ANC-HSS2010 and HTC2013 (R=0.6, p<0.001). All RPD sources and ANC-HSS indicated a decreasing trend in national HIV prevalence in Nigeria. PMTCT2014 data showed strong capability of predicting HIV prevalence in ANC-HSS2014 in regression model (B=2.09,p<0.0001). Use of routine PMTCT data in monitoring HIV prevalence among women of reproductive age could be more valid and reliable in local settings than the use of HTC data. Use of RPD to monitor national and sub-national-level HIV epidemic in between national surveys in Nigeria could maximize program resources, and promote a more responsive and efficient actions toward epidemic control.

## Introduction

An estimated 3.2 million people live with HIV (PLHIV) in Nigeria with 160,000 of those dying annually [1]. Additionally, the number of new infections was estimated at 220,000 in 2013 [1] which is the second highest in the African sub-region, while the disease has also rendered 1.8 million children orphans [2]. However, Nigeria has estimated a steady decline in the prevalence of HIV, from 5.8% in 2001, to 4.1% in 2010 and 3.0% in 2014 [3].

Common sources of HIV survey data are the Antenatal Care HIV Sero-Prevalence Sentinel Surveys (ANC-HSS) and the National AIDS and Reproductive Health Survey (NARHS). These surveys are designed for obtaining data for specific HIV related estimates that could be useful in population based health planning and programming. The ANC-HSS is conducted every 2 years [4] and NARHS every 4 years. ANC-HSS is conducted among all pregnant women who visit an antenatal facility for for anternatal services for the first time for a confirmed pregnancy [5]. The survey was conducted in about 160 antenatal care (ANC) sentinel facilities across the 36 states and the Federal Capital Territory (FCT)[6]. The NARHS is a nationally representative household survey conducted to elicit information from the general population on relevant HIV, reproductive health, and knowledge and behavioral indicators [7]. Though the NARHS is considered to be more nationally representative, it is more expensive and difficult to sustain in resource-limited settings.

Prevention-of-Mother-to-Child Transmission (PMTCT) of HIV services involves the identification of pregnant women livng with HIV during ANC and the provision of appropriate Antiretroviral Therapy (ART) to ensure that the mother stays healthy and prevent HIV transmission to the child [8]. The PMTCT services are provided in most ANC clinics so the women attending ANC services are easily referred to PMTCT services in the same clinic. Techically, routine PMTCT data and ANC-HSS both reflect HIV testing among pregnant women, however PMTCT data does not cover all ANC sites. Among pregnant women, enrolment into ART is preceded by an opt-out HIV test and counseling (HTC) process that determines the HIV status of the clients (including PMTCT clients). For the general population, HTC services are usually the entry points into HIV programs in which the clients are referred to access other HIV services (prevention, treatment, and care services) depending on their HIV status. Unlike the PMTCT services, which is limited to pregnant women, the HTC services are open to the general population in the clinics and communities through mobile community testing.

PMTCT data are primarily collected and generated at the HTC settings of the ANC clinics using the national PMTCT registers. The data are summarized on monthly basis by each facility using the Monthly Summary Forms (MSF) and transmitted manually or electronically to the next higher sub-national unit reporting/data collection level for validation, collation, and further transmittion to the national/central level. HTC data, sharing similar data reporting system with the PMTCT, is generated from all other HTC points of service operated at other settings within the clinic (such as voluntary counselling and testing, DOT-tuberculosis, out-patient-department settings etc) and community settings, except from the ANC; thus, its population is more generalized than the PMTCT and more likened to the NARHS population that involved household (community) testing.

Researchers have recommended the need to validate both the use of routine programmatic data (RPD) such as PMTCT program data for surveillance purposes [6,9–11] and the suitability of the ANC-HSS among other data sources in estimating the HIV prevalence in the general population [12–15]. These estimations form the essential principles of the Joint United Nations Programme on HIV/AIDS (UNAIDS) spectrum HIV estimates for many countries [16]. Some countries such as Kenya, Ethiopia [6] and Rwanda have been able to integrate the PMTCT information in the ANC forms which makes it easier to collect and compare information from both arms (RPD and Survey data) during the countries’ ANC-HSS. However, Nigeria has not been able to successfully link these two arms in ANC-HSS; hence, PMTCT data utility for surveillance and monitoring purposes has not been fully established in the country.

### Rationale for the study

The government of Nigeria (GoN) and PEPFAR have supported the generation and management of routine program data primarily for accountability and the monitoring of the PMTCT program, as well as to improve the implementation of HIV programs and services in Nigeria. However, the utility of program data in monitoring the country’s epidemic pattern and trend, and for timely impact assessment of the HIV programs in Nigeria has not been previously demonstrated. In additon, producing reliable, timely and consistent surveillance and population based data to support the effective implementation of evidence-based high-impact interventions to control HIV epidemic remains a challenge. The aim of the study was to determine the potential utility of RPD in monitoring HIV epidemic among the pregnant women and general population at the national and local levels in Nigeria. The significance of the study also shares in the benefits of effective HIV case base surveillance systems in which updated population level data that are routinely collected from health records are used to inform epidemic control decisions at both local and regional levels [17]; hence, accommodating the dynamism of the epidemic. The overarching research question was: to what extent could routine PMTCT and HTC program data sources be used to monitor HIV prevalence among women of reproductive age and in general population in the sub-national settings in Nigeria? This activity was reviewed in accordance with the US Centers for Disease Control and Prevention (CDC) human research protection procedures and was determined to be research, but did not involve human subjects.

### Objectives of the study

I. To determine data concurrence and any correlation(s) between HIV seropositivity estimates from RPD (PMTCT and HTC data) and prevalence estimates from the national survey/surveillance data (ANC-HSS and NARHS) sources between 2010 and 2014 in Nigeria.
II. To examine the trend and concurrence between the national HIV seropositivity rate from RPD and national prevalence from the national survey/surveillance data (NSD).
III. To determine the extent of predictive association between the HIV seropositivity estimates from RPD and prevalence from the NSD sources.

## Materials and methods

### Study design

The target population were the general population and pregnant women. We conducted a retrospective secondary analysis of HIV seropositivity data from RPD sources and compared them with those reported in NSD sources: ANC-HSS (for pregnant women) and NARHS survey (for general population) reports. The comparison, in principle, adopted the 12-step triangulation model recommended by WHO [18]. The model essentially involves initial identification of research questions that are answerable through triangulation, then, identifying the data sources, understanding the background of the data sources, collation of the data/reports, running observational analysis to understand the patern and trend among the sources, and drawing conculsions through summarized findings [18]. Data sources were identified based on the objectives of this study. The study variables from the data sources were the HIV prevalence estimates from NSD and seropositive rates from RPD.

### Data Sources

The relevant data sources were broadly categorized into two: NSD and RPD sources. The NSDs were the ANC-HSS and NARHS survey reports while the RPDs were the PEPFAR HTC and PMTCT APR data (see Table 1A below). The variables of interest from these data sources were the HIV prevalence estimates (from ANC-HSS and NARHS) and seropositivity rates (from PMTCT and HTC). The ANC-HSS and PMTCT data sources represent prevalence estimates among pregnant women while the NARHS and HTC data sources represent that in the general population.

**Table 1a.** Collated Data Sources for HIV Prevalence Estimates in Nigeria

| Data Categories | Data Sources | Years Available |  |  |  |  |
| --- | --- | --- | --- | --- | --- | --- |
|  |  | 2010 | 2011 | 2012 | 2013 | 2014 |
| NSD | ANC | Yes | N/A | No | N/A | Yes |
|  | NARHS | N/A | N/A | Yes | N/A | N/A |
| RPD | PEPFAR APR HTC | N/A | N/A | N/A | Yes | Yes |
|  | PEPFAR APR PMTCT | N/A | N/A | N/A | Yes | Yes |

*Note: No means that the data was expected for the year but not available or not in the format useful for the triangulation analyses. NA means the data was not expected for the year based on the original periodic design. None of the data sources were applicable (NA) in 2011, so the year was removed from the table.*

**Table 1b.**
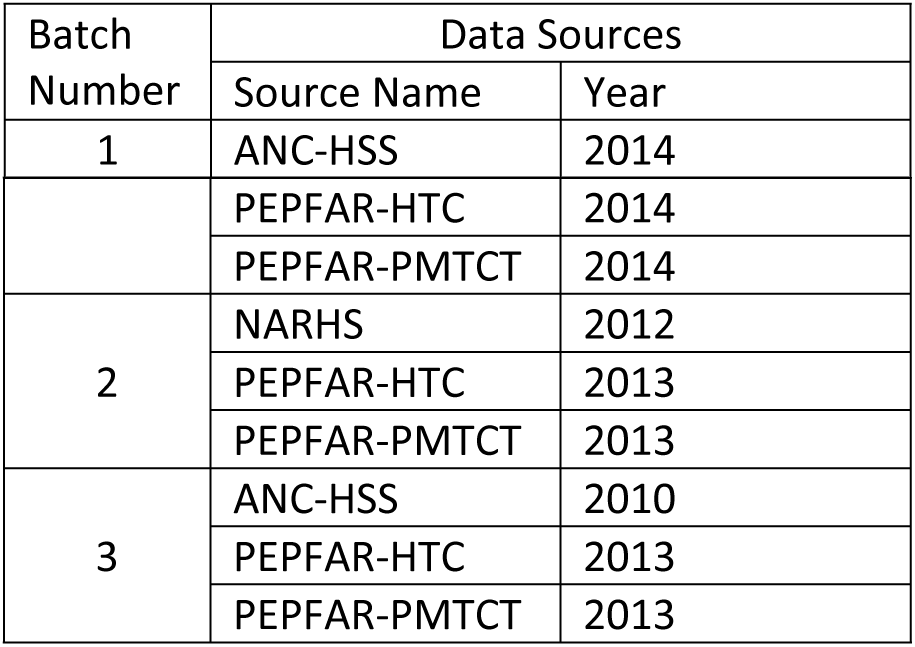
Batch grouping of the HIV Data Sources by year of availability for analyses

| Batch Number | Data Sources |  |
| --- | --- | --- |
|  | Source Name | Year |
| 1 | ANC-HSS | 2014 |
|  | PEPFAR-HTC | 2014 |
|  | PEPFAR-PMTCT | 2014 |
| 2 | NARHS | 2012 |
|  | PEPFAR-HTC | 2013 |
|  | PEPFAR-PMTCT | 2013 |
| 3 | ANC-HSS | 2010 |
|  | PEPFAR-HTC | 2013 |
|  | PEPFAR-PMTCT | 2013 |

National and states HIV prevalence and seropositivity estimates were abstracted from national reports and all the available data sources respectively, between 2010 and 2014. All avaibale PMTCT data (irrespective of contiguity of PMTCT facilties to ANC sentinel sites) were used to determine the state estimates. However, all four data sources of interest were not available for the same year due to the inconsistencies in the frequency at which national surveillance/surveys were conducted in Nigeria. The national surveys did not align with the years in which the RPD sources were available; thus, we paired the NSDs and the RPDs based on the closest years of availability (Table 1A).

Hence, the data sources were batched in the order of most matched or contiguous years and Batch 1 was most matched data sources (Table 1B).

### Data Analysis

Seropositive estimates were used as determined by the PEPFAR/HIV program. There were no advanced or further statistical adjustments to the methodologies or analysis. The program simply divided the number of HIV positive clients by the total number tested by states and any other sub national units (SNUs). Prevalence estimates from NSD sources were collated as reported in the national reports. We do not intend to manipulate any process or methods that yielded the estimates.

Visual analyses were performed to determine the geographic distribution patterns and concurrence levels of HIV prevalence and seropositivity using maps, and the level of correlation between the paired sources using the scatter plots charts. The scatter plot graphs were used to visualize the extent of correlation between the absolute prevalence estimates of the RPD sources and the NSD sources. A linear regression model line was fitted to show correlationship pattern as well as determine the correlation factor (R^2^) of the paired data sources. We used ArcMap 10.2 to generate visual maps and geographic distribution concurrence levels, all set at quantile classifications with four classes. The distribution concurrence levels were computed as the proportion of states (between the two compared data sources) that showed exactly the same HIV prevalence color patches. Technically, this means the number of states (between the paired sources in comparison) that fall within the same HIV prevalence quartile divided by the 37 (the 36 states plus Federal Capital Territory). These analyses were performed to respond to the first objective of the study.

Bivariate simple linear regression analysis were performed to determine the measures and strengths of the predictive relationships between the respective paired data sources in response to the third study objective. The SPSS-IBM 21.0 was used to perform the correlations and line charts, and linear regression modeling. For the linear regression, the RPD sources were used as the independent variables (predictors) and NSD sources as the dependent (response) variables, and leaving the entry method at “Enter”. This means that all the predictor/independent variables were entered into the model at once. Significance was assessed at P<0.05. Data sources from program and national surveys were analyzed in pairs to facilitate clarity within batches.

Trend analysis was performed on the respective national estimates (median of state-level estimates) from the data sources to determine the concurrence on their reflection of the trend of HIV prevalence in the country within the study period.

## Results

Analysis for Batch 1 data sources (ANC-HSS 2014, HTC 2014 and PMTCT 2014) showed that the median prevalence was lowest in the PMTCT 2014 (1.3%) and highest in the HTC 2014 (3.5%). The PMTCT data showed the most compact distribution (standard error[SE] 0.16), followed by HTC (SE 0.33) and ANC-HSS (SE 0.5). Batch 2 (NARHS 2012, PMTCT 2013, HTC 2013) showed similar pattern for HTC and PMTCT median prevalence, with HTC (4.2%) more than double that of PMTCT (1.6%). The NARHS 2012 survey had a median prevalence of 2.3%. Again, the PMTCT data showed a similar compact distribution (Standard deviation [SD] D 1.91) than the other data sources (NARHS: SD 3.34; HTC: SD 2.16) as with Batch 1 data sources. In Batch 3, median prevalence of ANC-HSS 2010 was 4.1%. These estimates stand between the lowest PMTCT 2013 and highest HTC 2013 median rates.

### Concurrence and Correlation Patterns

Figures 1-3 show line charts comparing states’ mean HIV prevalence rates between the data sources within each batch. Batch 1 graphs (Figure 1A&B) show that the ANC-HSS 2014 has a more correlated pattern with PMTCT 2014 at R^2^ = 44.0% than with the HTC 2014 at R^2^ = 23.3%

**Fig 1.Scatterplot of ANC-HSS 2014 and RPD sources 2014**

Figure 2, compares the mean HIV prevalence rates for the NARHS 2012, a National population based survey with PMTCT 2013 and HTC 2013 seropositivity rates. The NARHS 2012 source demonstrated low correlation patterns with the respective RPD sources at R^2^ less than 17%; however, it shows better correlation with HTC 2013 at R^2^ = 16.5% than with PMTCT 2013 at R^2^= 14.6%.

**Fig 2.Scatterplot of NARHS 2012 and RPD sources 2013**

Figure 3 shows that the correlation patterns of batch 3 paired data sources were similar to that of batch 2; however, with a stronger correlation factors between the paired ANC-HSS and RPD sources (R^2^ = 55.7% between ANC-HSS 2010 and PMTCT 2013 and R^2^ = 35.9% between ANC-HSS 2010 and HTC 2013).

**Fig 3.Scatterplot of ANC-HSS 2010 and RPD sources 2013**

#### Geographic Distribution Patterns of HIV Prevalence / Seropositivity rates

The geographic HIV prevalence patterns of the batch 1 data sources were similar (Fig 4). All show common higher HIV prevalence in the lower South-West, South-South, South-East, eastern part of the North-Central, and part of North-East regions of the country. ANC-HSS 2014 and PMTCT 2014 data from batch 1 had 40.5% (n=15)^1^ concurrence, and ANC-HSS 2014 and HTC 2014 data had 35.1% (n=13)^2^ concurrence. Generally, four states (Akwa-Ibom, Benue, Nassarawa, and Rivers States) had consistent prevalence patterns across the three data sources

**Fig 4.Map Geographic Distribution Pattern of HIV Prevalence from Batch 1 Data Sources.**

Batch 2 data sources showed a similar regional pattern as in batch 1 (see Fig 5). The distribution concurrence between NARHS 2012 and PMTCT 2013 was 40.5% (n = 15)^3^ and 37.8% (n=14) ^4^ for NARHS 2012 and HTC 2013.

**Fig 5.Map Geographic Distribution Pattern of HIV Prevalence from Batch 2 Data Sources.**

Comparing ANC-HSS 2010 to PMTCT 2013 and HTC 2013, the concurrence between ANC-HSS 2010 and PMTCT 2013 was 46% (n=17)^5^ and 37.8% (n=14)^6^ with HTC 2013 (Figure 6).

**Fig 6.Map Geographic Distribution Pattern of HIV Prevalence from Batch 3 Data Sources. Analysis of Correlations and Associations between the Data Sources**

In batch one, there was a significant correlation between mean HIV prevalence from PMTCT2014 program data and ANC-HSS 2014 surveillance report (R = 0.7, *p*< 0.01); and between the HTC2014 program HIV prevalence data and ANC-HSS 2014 surveillance report (R = 0.4, *p*< 0.05) as shown in table 3. The correlation between the HTC2014 program data and ANC-HSS 2014 data was statistically significant; however, it was rather weaker than the PMTCT2014 – ANC-HSS2014 correlation. In batch two, the individual correlations between the mean HIV prevalence of the two program data sources (PMTCT 2013 and HTC 2013) and the population based surveillance report (NARHS2012) were positive and statistically significant. PMTCT2013 and NARHS2012 was correlated as R = 0.38 (p< 0.05). HTC2013 and NARHS2012 was correlated at R = 0.41 (p< 0.05). The pattern of correlations among the paired sources in batch 3 were quite similar to that of batch 1. However, batch 3 correlations showed overall stronger statistically significant correlations between the 2013 RPD sources and ANC2010 report at the individual paired levels. The correlation between PMTCT2013 and ANC2010 was stronger (R = 0.75, *p*< 0.001) compared to that between HTC2013 and ANC2010 (R = 0.60, *p*< 0.001).

**Table 2.** Summary State Level HIV Prevalence Distribution from Routine and Surveillance Data Sources in Nigeria (2010-2014) n = 37

|  |  | ANC-HSS<br>2014 | PMTCT<br>2014 | HTC<br>2014 | NARHS<br>2012 | PMTCT<br>2013 | HTC<br>2013 | ANC-HSS<br>2010 |
| --- | --- | --- | --- | --- | --- | --- | --- | --- |
| Mean (SEM) |  | 4.00 (0.50) | 1.50 (0.16) | 3.91 (0.33) | 3.45 (0.55) | 2.28 (0.31) | 4.84 (0.35) | 4.80 (0.46) |
| Median |  | 3.0 | 1.28 | 3.47 | 2.30 | 1.62 | 4.20 | 4.10 |
| Std. Dev. |  | 3.03 | 0.96 | 2.01 | 3.34 | 1.91 | 2.16 | 2.80 |
| Min |  | 0.9 | 0.3 | 1.4 | 0.2 | 0.6 | 1.8 | 1.0 |
| Max |  | 15.4 | 4.7 | 10.4 | 15.2 | 9.2 | 11.0 | 12.7 |
| Perc. | 25 | 1.90 | .76 | 2.65 | 1.05 | 1.15 | 3.35 | 2.50 |
|  | 75 | 5.45 | 2.10 | 4.92 | 5.45 | 2.35 | 6.40 | 6.55 |

**Table 3.** Correlation and Regression Models of the paired RPD and PMTCT data sources

| Batch # | Dependent variable | Model (Predictors) | Pearson Correlation | $\beta$ beta coeff. (Std.Error) | R2 | 95% Confidence Interval for $\beta$ | |
| --- | --- | --- | --- | --- | --- | --- | --- |
|  |  |  |  |  |  | Lower Bound | Upper Bound |
| 1 | ANC-HSS 2014 | PMTCT2014 | 0.66*** | 2.09*** (0.39) | 43% | 1.28 | 2.90 |
|  |  | HTC2014 | 0.38** | 0.58* (0.24) | 12% | 0.10 | 1.06 |
| 2 | NARHS 2012 | PMTCT2013 | 0.38* | 0.67* (0.27) | 12% | 0.11 | 1.22 |
|  |  | HTC2013 | 0.41** | 0.63** (0.24) | 14% | 0.14 | 1.11 |
| 3 | ANC-HSS 2010 | PMTCT2013 | 0.75*** | 1.09*** (0.16) | 57% | 0.76 | 1.42 |
|  |  | HTC2013 | 0.60*** | 0.78*** (0.17) | 35% | 0.43 | 1.13 |

$n = 38$ . $p \leq 0.001$ \*\*\*, $p \leq 0.01$ \*\*, $p \leq 0.05$ \*

### Overall Comparative National HIV Prevalence Trend from Routine and Program Data Sources in Nigeria

On further review of the data sources to understand the HIV prevalence trend in the last three years among the three data sources with more than one time point for comparison (ANC-HSS, PMTCT and HTC), all suggested a steady decline in the HIV prevalence from 2012 to 2014 for PMTCT and HTC, and 2010 to 2014 for ANC-HSS (no middle year estimate was available) (Figure 7). However, the prevalence estimates from the HTC program data were consistently higher, while the prevalence estimates from the PMTCT program data were lower prevalence over the period. The ANC-HSS estimates, though interpolated on two extreme estimates, takes the middle values with HTC estimates providing the upper range and PMTCT providing the lower range along the trend.

**Fig 7.Trend Pattern of HIV Prevalence Estimates between 2010 and 2014 from the three data sources.**

### Predictive Association between Paired Data Sources

Table 3 also shows that the predictive association between PMTCT2014 program data and the ANC2014 HIV prevalence data in batch 1 was statistically significant at β = 2.09, (Confidence Interval [CI] 1.28, 2.90; R^2^=43%), and the association between the HTC 2014 and ANC-HSS 2014 prevalence estimates was also statistically significant at β = 0.6, (CI 0.10, 1.06; R^2^=12%). In batch 2, the association between NARHS2012 and the individual RPD sources were β = 0.67 (CI 0.11, 1.22; R^2^=12%) for PMTCT2013 and β = 0.63 (CI 0.14, 1.11; R^2^=14%) for HTC2013. In batch 3, the PMTCT2013 showed statistically significant association with ANC2010 at β = 1.09 (CI 0.76, 1.42; R^2^=57%) and HTC2013 showed similar association with ANC2010 but at β = 0.78 (CI 0.43, 1.13; R^2^=35%).

## Discussion

Our findings show that PMTCT data sources aligned more closely with the ANC-HSS data sources than HTC data sources did with the same ANC-HSS. This alignment pattern is understandable considering the fact that the ANC-HSS and PMTCT share similar population, which is pregnant women. On the other hand, though the PMTCT2013 and HTC2013 data demonstrated no appreciable correlation with the NARHS2012 source, and similar wideness to their fitted lines (R-squared < 17%), their patterns showed a positive relationships with the NARHS data. No clear differences in terms of individual visual alignments could be observed between the two RPD sources and NARHS; however, the ANC-HSS data seem to show a better visual alignment with PMTCT data sources than HTC sources.

The alignment pattern analysis of the state prevalence estimates in the different data batches were further supported by the comparative geographic distribution map analysis of the paired data sources. On comparing the geographic distribution pattern and concurrence levels of the RPD and NSD sources among the sub-national units, the PMTCT RPD and the corresponding NSD sources demonstrated higher concurrence levels across the three batches than seen with the HTC RPD sources. For example, the PMTCT RPD source concurrence level from batch 1 to 3 were 40.5%, 40.5%, and 46% compared to 35.1%, 37.8%, and 37.8% with HTC RPD sources. The map review, in addition, showed similarities in the general regional distribution pattern in HIV prevalence estimates between the paired data sources. They demonstrated strong evidence that the higher burden of the disease may reside at the South-Eastern, Central-Eastern, and extreme North-Eastern Nigeria. This further demonstrated that the PMTCT data could provide similar regional estimates of the disease burden than the HTC data with reference to the ANC HIV sero-prevalence sentinel survey estimates. In all, the map analysis strengthens the similarities and utility of RPD in monitoring the country prevalence using four classification levels of the prevalence.

In all instances, PMTCT estimates had stronger significant association with both ANC-HSS and NARHS estimates compared with HTC, but more with ANC-HSS than NARHS. This could be explained by the common target population from which the ANC-HSS and PMTCT programmatic data sources were generated, and similar in methods and settings. These findings were consistent with the findings observed in similar studies in India, [19] Rwanda, [10] and Mozambique [11]. However, the utility of PMTCT data in substituting ANC-HSS data had been questioned, largely due to perceived poor quality in the testing and data management practices at PMTCT program sites [12-14]. Bolu et al., however, suggested that despite these reservations, the PMTCT still has better utility in HIV program management when used in conjunction with the ANC-HSS data [6].

Our findings demonstrated a decreasing trend in the national seropositivity estimates from 2012 to 2014. This trend is similar to that of the national HIV prevalence observed in ANC-HSS source from 2010 to 2014; however, only two estimates in 2010 and 2014 were available. The trend alignment is explained by the higher to lower mean seropositivity rate / prevalence derived from these sources. The trend from NARHS could not be ascertained because only one prevalence estimate in 2012 was available. Furthermore, the relatively low prevalence estimates from the PMTCT data sources (compared to ANC-HSS) in this study were similar to those reported in the literature. Earlier studies found that the PMTCT prevalence data were general lower than that of the ANC-HSS [6,10,11] except Seguy et al. who found the PMTCT prevalence (14%) a little higher than the ANC-HSS prevalence (13%) [9]. Our results suggested that PMTCT prevalence estimates may be lower than the national survey prevalence while the HTC prevalence estimates are higher than the national surveys; thus, PMTCT prevalence may form the lower range of the national surveys while the HTC estimates forms the upper range limits.

Although our study data sources did not include sufficient population-based survey data, the results concur with the spectrum suggested that the ANC-HSS may overestimate the prevalence in the general population but could still reliably estimate the HIV prevalence trend in the general population. The suggestion is consistent with the conclusions from similar comparative studies in 26 countries with generalized epidemics, [12] in Zimbabwe, [13] in sub-Saharan Africa [14] and in Uganda [15] in which the ANC-HSS estimated prevalence were slightly higher than the prevalence observed in the population-based surveys. In their detailed review of the correlations, Gouws et al. suggested that the ANC-HSS data be multiplied by 0.8 to adjust for epidemic trend in the general population [12]. There is an essential difference between the NARHS and ANC-HSSs, either in their respective primary target populations or the methodology or both.

The HTC data is generated from across the entire population groups by age and sex but only from those who visited the health and mobile community testing facilities, unlike the NARHS data that is generated across the entire population groups from the households and through scientific sampling. This could explain the wide differences in the similarity of the program data with the NARHS, however, in all, NARHS data showed a slightly closer data fitting with the HTC than the PMTCT data. We observed that the HTC data overestimated the ANC-HSS prevalence, in contrast to the under-estimation of the PMTCT data. The declining trend pattern observed across the ANC, HTC, PMTCT prevalence estimates further shows that the HIV epidemic in Nigeria may be on the decrease. The three data sets all agree on this pattern and further strengthen the potential use of the raw programmatic data for monitoring of the country’s HIV epidemic trend.

The observed correlations agree with our modelled results to a large extent. Our finding suggests that PMTCT program data could reflect or closely monitor state level epidemic pattern obtainable with the ANC-HSS data, but may not produce exact point estimates. Our findings suggest that the PMTCT seropositivity may be multiplied by 2.0 to adjust for ANC-HSS prevalence estimates. Generally, the findings agree with that of most previous related studies, pulling more weight to the usefulness of PMTCT data. This further reveals the potential direction of research toward the exploration of the use of other RPD from other indicators and disease programs for local health surveillance. Such future studies could be a break-through for less-expensive, easy, and fairly reliable m for health surveillance in resource-limited countries.

### Limitations

This study is subject to at least four limitations. First, except for batch 1 data sources, other batch data sources were not from the same year; thus, their prevalence estimates may not be a perfect match. Secondly, the RPD were generated from the health facilities across the country; however, the facility distribution pattern of the country was not determined in this study, hence, could not determine the representativeness of the RPD to inform the appropriateness of estimating population based surveys such as NARHS and DHS [7,20]. The implication of this is that the potential utility of the RPD for monitoring national HIV epidemic pattern may change with change in the distribution of the facilities that generate these data. These findings assume that facilities were fairly distributed all over the country. In addition, the quality of the respective methodologies, HIV testing services in the facilities, and data managements of the data sources could not be determined in this study and may improve the correlation analyses of the data sources. Lastly, the concurrence analysis that quantified the geographic distribution accounted for the exact concurrence. This means that a state with level 4 and 3 prevalence between two data sources were considered as non-concurrence in the analysis regardless of the closeness. Such closeness was considered same as level 4 and 1 between the data sources. However, direct visual review of the maps could help resolve this limitation.

## Conclusions

The study findings suggest that the RPD sources could be significantly correlated and associated with the NSD sources in monitoring the HIV prevalence in the country. The RPD sources could reliably demonstrate the trend in HIV epidemic at national and state levels. As expected, PMTCT data tend to underestimate the ANC-HSS HIV prevalence while the HTC data showed indication of overestimation of the epidemic trend among the reproductive female population. The potential use of routine PMTCT programmatic data may be more reliable and favorable than HTC program data in estimating the national HIV epidemic among women of reproductive age in Nigeria. The implication of the finding is that, PMTCT could largely be a more promising source for HIV surveillance in the local settings in Nigeria and other resource-limited countries, particularly in settings where no monitoring information exists or periodic population-based surveys are infrequent.

## Acknowledgements

This research [or project, publication, etc.] has been supported by the President’s Emergency Plan for AIDS Relief (PEPFAR) through the Centers for Disease Control and Prevention (CDC)

The authors acknowledge and appreciate the thorough reviews and edits made by Dr. Stacie Greby.

## Supporting information

**S1 Fig. Scatterplot of ANC-HSS 2014 and RPD sources 2014**

**S2 Fig. Scatterplot of NARHS 2012 and RPD sources 2013**

**S3 Fig. Scatterplot of ANC-HSS 2010 and RPD sources 2013**

**S4 Fig. Map Geographic Distribution Pattern of HIV Prevalence from Batch 1 Data Sources**

**S5 Fig. Map Geographic Distribution Pattern of HIV Prevalence from Batch 2 Data Sources**

**S6 Fig. Map Geographic Distribution Pattern of HIV Prevalence from Batch 3 Data Sources**

**S7 Fig. Trend Pattern of HIV Prevalence Estimates between 2010 and 2014 from the three data sources.**

**S1 Table. Study dataset**

## Footnotes

1 ANC-HSS 2014:PMTCT 2014 states with similar pattern (Kebbi, Zamfara, Jigawa, Yobe, Oyo, Ekiti, Bayelsa, Rivers, Akwa Ibom, Ebonyi, Benue, Nasarawa, FCT, Enugu and Adamawa).

2 ANC-HSS 2014:HTC 2014 states with similar pattern (Kebbi, Zamfara, Jigawa, Bauchi, Oyo, Edo, Delta, Bayelsa, Rivers, Akwa Ibom, Benue, Nasarawa and Ogun).

3 NARHS 2012:PMTCT 2013 states with similar prevalence patterns (FCT, Nassaraw, Akwa Ibom, Rivers, Taraba, Cross-River, Osun, Jigawa, Kwara, Enugu, Bauchi, Katsina, Zamfara, Kebbi, and Ebonyi).

4 NARHS 2012:HTC 2013 states with similar prevalence patterns (Katsina, Zamfara, Kebbi, Osun, Ekiti, Edo, Kogi, Enugu, Enonyi, Cross River, Rivers, Akwa Ibom, Taraba, and Plateau)

5 ANC-HSS 2010:PMTCT 2013 states with similar prevalence patterns (FCT, Nassaraw, Benue, Akwa Ibom, Rivers, Abia, Borno, Kogi, Lagos, Niger, Oyo, Ogun, Yobe, Bauchi, Katsina, Zamfara, and Kebbi)

6 ANC-HSS 2010:HTC 2013 states with similar prevalence patterns (Benue, Akwa Ibom, Abia, Bayelsa, Kaduna, Lagos, Niger, Ogun, Delta, Katsina, Zamfara, Kebbi, Kwara, and Ekiti)

